# Osprey: Open-Source Processing, Reconstruction & Estimation of Magnetic Resonance Spectroscopy Data

**DOI:** 10.1101/2020.02.12.944207

**Authors:** Georg Oeltzschner, Helge J. Zöllner, Steve C. N. Hui, Mark Mikkelsen, Muhammad G. Saleh, Sofie Tapper, Richard A. E. Edden

**Author notes:** **Corresponding author:** Georg Oeltzschner, Ph.D., Division of Neuroradiology, Park 367G, The Johns Hopkins University School of Medicine, 600 N Wolfe St, Baltimore, MD 21287.

## Abstract

**Background:** Processing and quantitative analysis of magnetic resonance spectroscopy (MRS) data are far from standardized and require interfacing with third-party software. Here, we present Osprey, a fully integrated open-source data analysis pipeline for MRS data, with seamless integration of pre-processing, linear-combination modelling, quantification, and data visualization.

**New Method:** Osprey loads multiple common MRS data formats, performs phased-array coil combination, frequency-and phase-correction of individual transients, signal averaging and Fourier transformation. Linear combination modelling of the processed spectrum is carried out using simulated basis sets and a spline baseline. The MRS voxel is coregistered to an anatomical image, which is segmented for tissue correction and quantification is performed based upon modelling parameters and tissue segmentation. The results of each analysis step are visualized in the Osprey GUI. The analysis pipeline is demonstrated in 12 PRESS, 11 MEGA-PRESS, and 8 HERMES datasets acquired in healthy subjects.

**Results:** Osprey successfully loads, processes, models, and quantifies MRS data acquired with a variety of conventional and spectral editing techniques.

**Comparison with Existing Method(s):** Osprey is the first MRS software to combine uniform pre-processing, linear-combination modelling, tissue correction and quantification into a coherent ecosystem. Compared to existing compiled, often closed-source modelling software, Osprey’s open-source code philosophy allows researchers to integrate state-of-the-art data processing and modelling routines, and potentially converge towards standardization of analysis.

**Conclusions:** Osprey combines robust, peer-reviewed data processing methods into a modular workflow that is easily augmented by community developers, allowing the rapid implementation of new methods.

## 1. Introduction

Magnetic resonance spectroscopy (MRS) is the only methodology that can determine the levels of neurochemicals in living tissue non-invasively, providing a unique window on the neurobiology of the human brain in health and pathology. Over the course of several decades, the field has developed a wide range of data acquisition, processing, and quantitative analysis methods (Harris et al., 2017; Landheer et al., 2019; Öz et al., 2020; Wilson et al., 2019).

In general, state-of-the-art analysis of MRS data can be divided into three fundamental steps:

1. **Pre-processing** of raw data that has been exported directly from the scanner or obtained from an institutional archiving system (PACS). Currently, no convention for a standardized MRS data format exists. Instead, each major vendor has developed proprietary file formats that store data in varying degrees of ‘rawness’ and limited information on acquisition parameters. Some of the most widely used file formats (DICOM MRS, Philips SDAT/SPAR, Siemens RDA, GE P-file) contain data that have been pre-processed to varying extents in the vendor-native online reconstruction environment. On-scanner processing relies on proprietary vendor-specific reconstruction code and is therefore neither standardized nor publicly documented. Data processing can include basic low-level operations on the raw time-domain data (down-sampling, zero-filling, filtering, truncating, Fourier transformation), higher-level operations to improve critical signal properties like linewidth and signal-to-noise ratio (weighted receiver-coil combination, alignment of individual averages), and operations to address acquisition-related artefacts (removal of residual water signal, eddy-current correction).
2. **Modelling** of the processed spectral data is performed to derive quantitative parameters that allow conclusions to be drawn about the levels of individual metabolites. The complexity of this process ranges from simple peak integration, through single-resonance modelling, to linear-combination algorithms to decompose spectra into their constituent signals. While simple models are easy to implement, the full information content of an MRS spectrum can only be unlocked by modelling it fully. The most widely used linear-combination algorithms are exclusively implemented in third-party compiled software, either open-source (e.g. Tarquin (Wilson et al., 2011), Vespa (Soher et al., 2011)), or closed-source academic (e.g. the AQSES (Poullet et al., 2007) algorithm in jMRUI (Stefan et al., 2009)) or commercial (LCModel (Provencher, 1993)). Despite their widespread use, all common fitting software packages are developed and maintained by small groups of researchers (or even a single individual), who often critically rely on third-party funding to keep the project alive.
3. **Quantification** is here used to describe the process of converting quantitative modelling parameters into biologically meaningful estimates of metabolite levels. Depending on the complexity of the quantification method, this may entail simply taking ratios of peak areas, or include more sophisticated calculations such as correcting for the fraction of cerebrospinal fluid (CSF), tissue-specific relaxation correction (Gasparovic et al., 2006), or accounting for assumed differences of metabolite abundance between tissue classes (Harris et al., 2015).

The core task of *modelling* is usually performed by third-party software, which typically has limited capability of *pre-processing* and *quantification*. This forces researchers to create their own local pipeline, starting from a rich diversity of scanner-specific export formats, choosing an appropriate set of processing steps, and finally exporting the processed data into a format that is accepted by the modelling software. Many modelling software solutions include the calculation of signal amplitude ratios to a reference (creatine, N-acetyl aspartate, water), but at the time of writing, none of them allow direct incorporation of tissue-specific segmentation or metabolite-specific relaxation information. Therefore, researchers must, again, develop local custom code to import the modelling results, apply corrections, and calculate final quantitative measures. In contrast to popular imaging modalities like functional MRI, which have established publicly available analysis frameworks and software environments open to community contributions, a widely used standardized pipeline is currently not available for MRS data analysis.

We have identified several issues with these practices. In short, the current best practices are not only inefficient, but severely hinder the more widespread use of MRS as a research tool, and curb its potential as a clinical one:

a. **Waste of resources:** If every lab resorts to writing custom code to carry out the same task, a lot of time, energy, and funding is wasted into duplicate efforts.
b. **Methodological heterogeneity:** The lack of a single analysis pipeline to include pre-processing, modelling and quantification forces all labs to rely on local custom code. Additionally, many labs conducting advanced methodological MRS research rely on their own, long-established pipelines (including either local or third-party modelling) and keep methodological developments local. This is detrimental to standardization, comparability, and transparency of data analysis.
c. **High-entry threshold:** Developing custom analysis code represents an often insurmountable effort for new research groups for whom MRS is a potential tool, but not a primary focus, and who might not have the background knowledge, resources, or technical expertise to create a processing pipeline from scratch.
d. **Inertia and slow evolution:** New methodological developments take longer than necessary to gain acceptance and become widely adopted, because newcomers struggle to implement them if the code is not made publicly available right away. Integrating new developments into compiled third-party software requires larger programming efforts, with low incentives for the developer to devote resources to this task.
e. **Dependence and vulnerability:** To evolve and be maintained, third-party software is critically dependent on its developers. They may transition to different positions (or leave academia altogether), run out of funding, lack time or staff support, or simply have no incentive to actively develop the software. This is particularly true for closed-source or commercial software, or tools that are maintained by single individuals.

Here we describe a new open-source MATLAB-based toolbox “Osprey”. Osprey is an all-in-one software package that combines all steps of state-of-the-art pre-processing, linear-combination modelling, quantification, and visualization of MRS data. The Osprey framework is designed as a modular, fully open-source environment to flexibly adopt future methodological developments, accelerate their adaptation, and foster standardization. The entire source code of Osprey is available in the public domain, inviting improvements, bug fixes, and addition of state-of-the-art technical developments from the community.

## 2. Methods

The Osprey workflow is summarized in Fig. 1. Osprey consists of seven separate modules – ***Job***, ***Load***, ***Proc***, ***Fit***, ***Coreg***, ***Seg***, and ***Quant***, all of which are sequentially called by simple MATLAB commands. Alternatively, users can conduct the entire workflow in a graphical user interface designed to minimize the amount of user input. The Osprey code builds upon functions and the organizational structure of FID-A (Simpson et al., 2017), an open-source collection of MATLAB scripts for simulating, loading, and processing MRS data. At the time of writing, the various signal processing and modeling functions require the Image Processing Toolbox (to handle DICOM files), the Optimization Toolbox (to model the spectra), the Signal Processing Toolbox (for Hilbert transforms), and the Statistics and Machine Learning Toolbox (for statistical analysis). To use the Osprey GUI, the GUI Layout Toolbox (Sampson, 2019) and Widgets Toolbox (Jackey, 2019) need to be installed from the MATLAB File Exchange. Osprey has been tested using MATLAB versions 2017a and newer.

**Figure 1:**
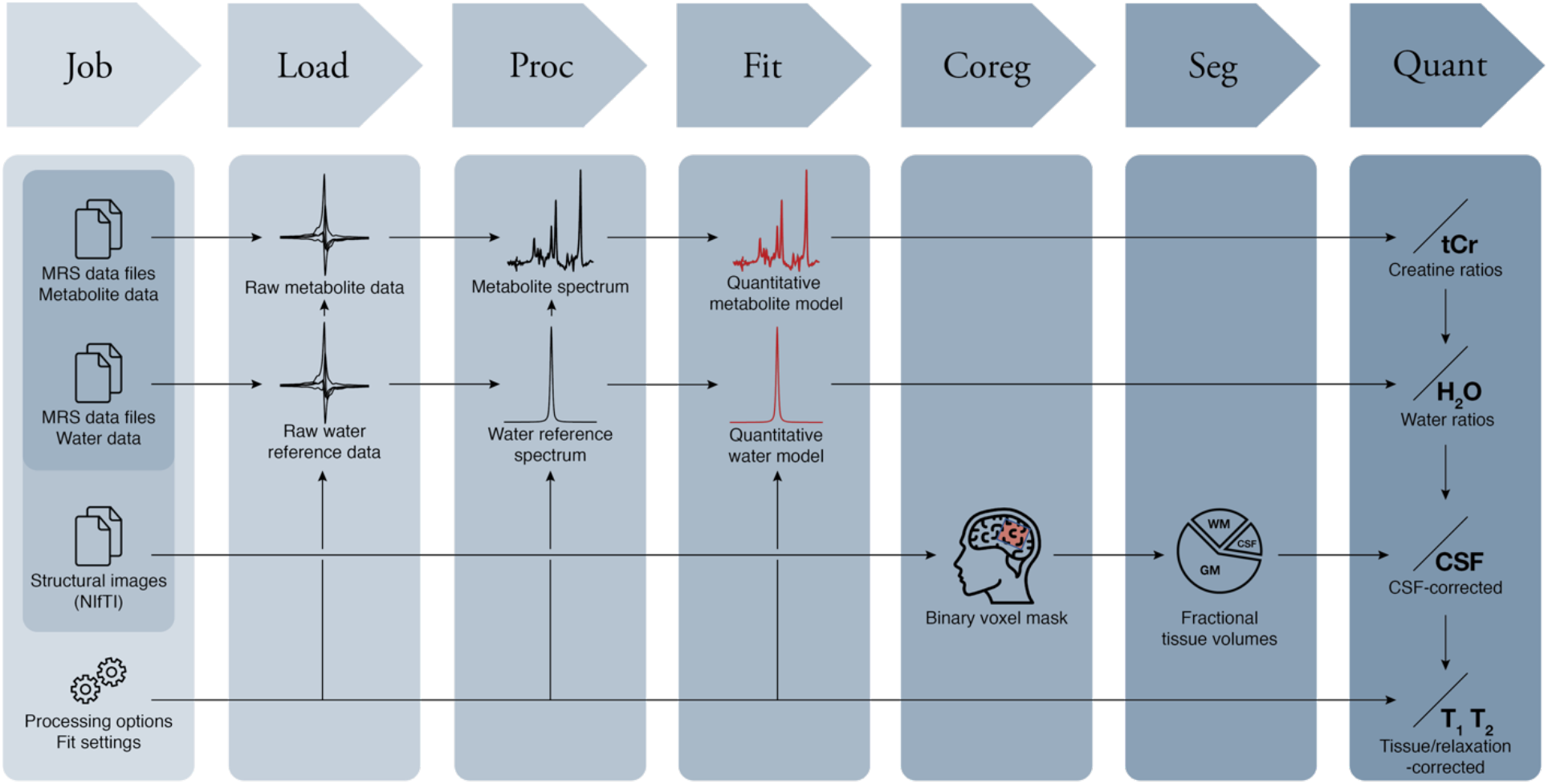
The Osprey workflow with the seven modules Job, Load, Proc(ess), Fit, Coreg(ister), Seg(ment), and Quant(ify).

Additional details and MATLAB commands are summarized in section S1 of the Supplementary Material.

### 2.1 Job

The only user interaction that is required to specify and conduct an Osprey data analysis is to define a ***Job*** in a job file. The job system allows easy batch processing of multiple datasets. The job file is a text file containing the paths to the MRS data files and structural image files to be processed, and parameters to control data processing and linear-combination modelling. In addition, the user must specify an output folder that Osprey will use to save exported data, coregistration and segmentation images, and quantitative result tables.

Osprey distinguishes between three classes of MRS data files:

- Metabolite (water-suppressed) data. These are a mandatory input.
- Lineshape reference data. These are an optional input, acquired with the same sequence as the metabolite data, but without water suppression, and used to perform eddy-current correction (Klose, 1990) of the metabolite data. If only lineshape reference data are provided, this signal is also used to calculate water-scaled concentration estimates.
- Short-TE reference data. These are another optional input. If the user provides short-TE reference data, they will be used to derive water-scaled concentration estimates (and lineshape data are only used for eddy-current correction). Using short-TE water as the concentration reference standard reduces T2-weighting of the water reference signal (and associated correction errors) compared to long-TE water data.

In addition to the paths to the raw data files, the job file must specify the type of sequence (with or without spectral editing, such as MEGA (Mescher et al., 1998; Rothman et al., 1993), HERMES (Chan et al., 2016; Saleh et al., 2016), or HERCULES (Oeltzschner et al., 2019a)), the target molecules of spectral editing experiments, and options for the fitting process, which are explained in Section 2.4.

### 2.2 Load

Osprey supports most common raw and processed file formats from major MRI vendors. This includes Philips (SDAT/SPAR, DATA/LIST), Siemens (RDA, TWIX) and GE (P-file) data. Additionally, single-file or multi-file DICOM datasets can be loaded. At the time of writing, single-voxel conventional and various *J*-difference-edited data from many sequence implementations are supported.

Upon calling the ***Load*** command, Osprey determines the file format, loads the raw spectroscopic data (FIDs) as well as all header information necessary for subsequent modules. The *Load* module also performs receiver-coil combination weighted with the ratio of the signal to the square of the noise. If lineshape reference or short-TE water data are provided, they are used instead of the metabolite data to determine the phasing and weighting parameters.

### 2.3 Proc

The ***Proc*** (Process) module performs all necessary steps to translate the raw, un-aligned, un-averaged data into spectra that are ready to be modeled.

The pre-processing pipeline includes the following steps: 1) alignment of individual averages using spectral registration in the time domain (Near et al., 2015); 2) averaging; 3) Fourier transformation; 4) automatic determination of the correct polarity of the spectrum since some water-suppression schemes can result in negative residual water peaks; 5) residual water removal by singular-value decomposition of the signal (Barkhuijsen et al., 1987) and subtracting components between 4.6 and 4.8 ppm; 6) linear baseline correction (based on the mean of 100 data points at the far edges of the frequency domain spectrum; 7) correct frequency referencing based on a single-Lorentzian fit to the 2.01 ppm NAA singlet. If lineshape reference data are available, the Klose method (Klose, 1990) is applied to correct the metabolite spectra for eddy-currents.

For *J*-difference-edited experiments (MEGA, HERMES, HERCULES), sub-spectra are aligned by minimizing the frequency-domain difference signal within a particular frequency range containing identical signal in pairs of sub-spectra. As an example, the edit-ON and edit-OFF spectra in GABA-edited MEGA data usually share a considerable residual water signal that is used for alignment. In contrast, the residual water signal is suppressed in the edit-ON spectra of GSH-edited MEGA data due to the editing pulse being applied at 4.56 ppm. In this case, the 2.01 ppm NAA signal is used for alignment. After sub-spectrum alignment, difference and sum spectra are calculated and stored.

In addition to performing the automated processing, Osprey determines several quality-control parameters. Linewidth is determined as the full-width half-maximum of a single-Lorentzian fit to the NAA peak (between 1.8 and 2.2 ppm). SNR is determined as the ratio between the amplitude of the NAA peak and the standard deviation of the detrended noise between −2 and 0 ppm. The frequency drift over the course of the experiment is determined based on the creatine signal in every single average (creatine and choline signals that nominally appear at 3.02 and 3.20 ppm are modeled by two Lorentzians).

The *Proc* module can optionally export the fully processed metabolite and water reference spectra to output subfolders, in formats readable by external third-party fitting software (LCModel, Tarquin, jMRUI, Vespa). This feature allows users to perform traditional data modelling, with the benefit of improved SNR and linewidth resulting from optimized coil-combination and alignment of individual averages – features that the established software solutions currently do not offer.

### 2.4 Fit

The ***Fit*** module models the processed spectra passed from the *Proc* module with a linear combination of basis functions. The default Osprey model is designed to mimic several key features of the algorithms implemented in LCModel and Tarquin. Several fit options are specified in the job file, e.g. the frequency range over which the spectrum is to be modelled, and the baseline flexibility (as controlled by the minimum ppm-spacing between neighboring knots of the cubic baseline spline).

The *Fit* module requires a sequence-specific basis set, which is automatically selected by Osprey based on the vendor, sequence type, and sequence parameters. Several sequence-specific basis sets for commonly used implementations of PRESS, MEGA-PRESS, HERMES, and HERCULES are included, derived from fast spatially resolved density-matrix simulations (Zhang et al., 2017) using ideal excitation and shaped refocusing pulses. Basis functions for macromolecule and lipid functions are added by generating Gauss-shaped signals with properties summarized in Tables S1 and S2 in section S3 of the Supplementary Material.

The *Fit* module interpolates the basis set to match the resolution (data points per ppm) of the processed spectra. All spectra of a single dataset (water-suppressed and water-unsuppressed) are then scaled to the basis set to facilitate convergence of the subsequent optimization.

Prior to the full analysis, two preparation steps are carried out. First, for optimal frequency referencing, the spectrum is cross-correlated with a sum of unit delta functions at 2.01 ppm, 3.03 ppm, and 3.21 ppm, representing the major landmark singlets from NAA, Cr, and Cho, respectively, and the frequency shift corresponding to the offset of the cross-correlation function is applied to the spectrum. Second, to obtain good starting values for the phase and linebroadening parameters, a preliminary fit is performed with a reduced basis set only including the basis functions of NAA, Cr, PCh, Glu, and Ins, and a more flexible baseline with a knot spacing of 0.15 ppm. The final phase and linebroadening estimates from this preliminary fit are used as starting values for the full fit. Together with the initial referencing step, selecting a reasonable starting point for the non-linear parameters helps stabilize the optimization problem.

Osprey fits the real part of the frequency-domain spectrum *Y*(*v*) using a model similar to the one used by the LCModel algorithm. The *N*_*M*_ simulated time-domain metabolite basis functions *m*_*m*_(*t*) in the basis set receive the same Gaussian linebroadening *γ*, and individual Lorentzian linebroadenings *α*_*m*_ and frequency shifts *ω*_*m*_ (*m* = 1, …, *N*_*M*_) before they are Fourier-transformed into the frequency domain:

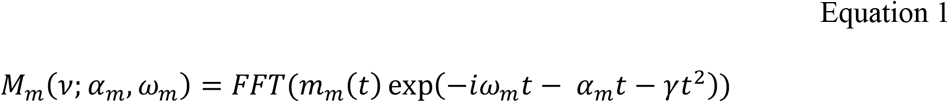

To account for deviations from a perfect Voigtian lineshape as determined by the linebroadening parameters, the frequency-domain basis spectra are then convolved with an arbitrary, unregularized lineshape model. This supplementary lineshape model ***S*** has a length equal to a spectral width of 2.5 times the coarse estimate of the FWHM of the spectrum that was estimated during the initial referencing step. ***S*** is normalized, so that this convolution does not impact the integral of signals, and is initialized as a unit delta function at the central point.

The smooth baseline is constructed as a linear combination of *N*_*B*_ normalized, equally spaced cubic B-spline basis functions *B*_*j*_(*v*) with coefficients *β*_*j*_ (*j* = 1, …, *N*_*B*_), with knots 2 and (*N*_*B*_ − 1) located on the edges of the fit range, and two additional knots outside the modeled range. *N*_*B*_ is determined from the fit range based upon the minimum knot spacing condition that the user specifies in the job file. In contrast to the LCModel algorithm, the default Osprey model does not currently include baseline regularization. To prevent an unreasonably flexible baseline without using a regularizer, the default Osprey spline knot spacing is increased to 0.4 ppm, compared to the default LCModel ‘DKNTMN’ setting of 0.15 ppm.

The model spectrum 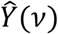 is constructed from these components as follows:

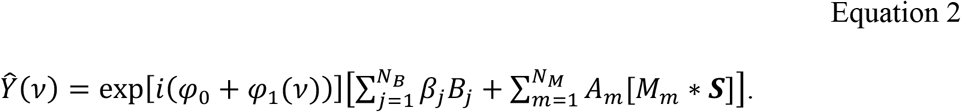

Here, *φ*_0_ represents the global zero-order phase correction; *φ*_1_(*v*) the global first-order (linear) phase correction; and *A*_*m*_ the amplitude of each metabolite/MM/lipid basis function.

To minimize the sum of squares of the difference between the data *Y*(*v*) and the model 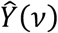, the *Fit* module uses an implementation of the popular Levenberg-Marquardt (Levenberg, 1944; Marquardt, 1963) non-linear least-squares optimization algorithm that allows hard constraints to be imposed on the parameters (Dentler, 2015). The amplitude parameters applied to the metabolite and baseline spline basis functions occur linearly in the model, and are determined at each iteration of the non-linear algorithm with a limited-memory algorithm for bound constrained optimization (L-BFGS-B) (Becker, 2015; Byrd et al., 1995; Zhu et al., 1997), constraining the metabolite amplitudes to be non-negative (*A*_*i*_ ≥ 0). Default hard constraints on non-linear parameters and weak soft constraints on macromolecule and lipid amplitudes are imposed to stabilize the solution, and are defined as they are in LCModel and Tarquin (Table S3 in section S4 of the Supplementary Material).

Water-unsuppressed data are modelled from a simulated water resonance with the same constrained non-linear least-squares algorithm using a simplified model, only including the following six hard-constrained modelling parameters: zero-order phase (−2*π* ≤ *φ*_0_ ≤ 2*π*), first-order phase 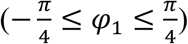, Gaussian linebroadening 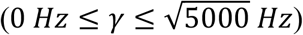, Lorentzian linebroadening (0 *Hz* ≤ *α* ≤ 50 *Hz*), frequency shift (−15 *Hz* ≤ *ω* ≤ 15 *Hz*), and amplitude *A*. No baseline is included in the water model.

### 2.5 Coreg

The ***Coreg*** module uses information about size, position and orientation of the MRS voxel in scanner-space coordinates to create a binary voxel mask, i.e., a 3D image in which the values 1 and 0 represent locations inside or outside the MRS voxel, respectively. The binary voxel mask is then transformed to the same coordinate system as the structural image (in NIfTI format) that the user provides in the job file. This step ensures that the voxel mask is coregistered to the structural image, and reproduces the original voxel placement.

### 2.6 Seg

The ***Seg*** module invokes the SPM12 segmentation function to segment the structural image into gray matter (GM), white matter (WM), and cerebrospinal fluid (CSF). The coregistered voxel mask that was created by the *Coreg* module is then overlaid with the GM, WM, and CSF tissue probability maps. The ***Seg*** module then calculates fractional tissue volumes *f*_*vol*_ for GM, WM and CSF according to:

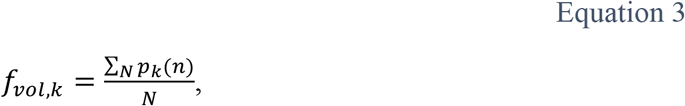

with the tissue probabilities *p*_*k*_(*n*) for the tissue class *k* ∈ *GM*, *WM*, *CSF* and the *n*-th image voxel (*n* ∈ 1, 2, …, *N*), where *N* represents the number of image voxels within the MRS voxel.

### 2.7 Quant

The ***Quant*** (Quantify) module calculates various quantitative outputs, depending on the available modelling parameters that have been determined during the *Fit* process:

- Ratios of the metabolite signal amplitudes *S*_*met*_ to the total creatine amplitude *S*_*tCr*_ = *S*_*Cr*_ + *S*_*PCr*_ are always determined, regardless of whether water data have been provided, according to

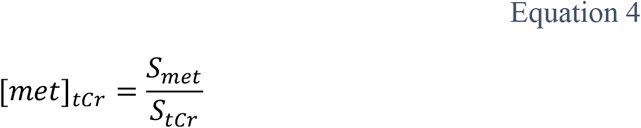

These tCr ratios are reported as raw ratios, i.e. no relaxation correction or accounting for tissue composition is applied. In difference-edited experiments, the Cr resonances are usually subtracted out in the difference spectra. In these cases, the tCr reference is determined from the edit-OFF spectrum if the fit option ‘Separate’ has been selected, and from the sum spectrum if the fit option ‘Concatenated’ has been selected.
- When an unsuppressed water signal is provided, Osprey can report water-scaled metabolite estimates. If both lineshape reference data (i.e. data with the same TE as the water-suppressed data) and additional short-TE water data are available, the latter will be used as the water-scaling reference signal. Osprey reports water-scaled metabolite estimates according to Equation 5, analogous to the LCModel water scaling procedure, which does not account for tissue composition and assumes pure white matter by default:

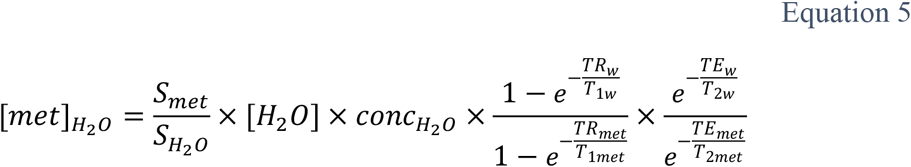

Here, [*H*_2_*O*] is the molal concentration of pure MR-visible water, i.e. 55.5 mol/kg of MR-visible water (Gasparovic et al., 2006; Knight-Scott et al., 2003); 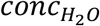 is the relative water density of white matter (0.65); *TR*_*w*_, *TR*_*met*_, *TE*_*w*_, *TE*_*met*_ are the repetition and echo times of the water-unsuppressed and water-suppressed acquisitions; *T*_1*w*_ and *T*_2*w*_ are averaged relaxation times for tissue water (for brain data at 3T, *T*_1*w*_ = 1100 ms and *T*_2*w*_= 95 ms (Wansapura et al., 1999)); *T*_1*met*_ and *T*_2*met*_ are the averaged relaxation times of all metabolites and generated from a lookup table that can be modified by the user. No tissue correction is applied. These water-scaled estimates are calculated even if no tissue composition information (from the *Seg* module) is available.
- When unsuppressed water data and tissue segmentation are available, Osprey calculates water-scaled metabolite estimates corrected for the volume fraction of CSF in the voxel according to

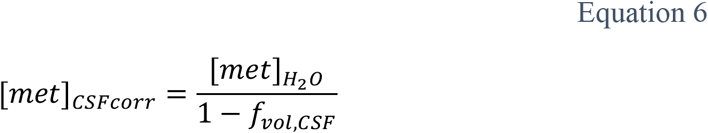

Here, 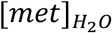 is the water-scaled metabolite estimate obtained in the previous step, and *f*_*vol*,*CSF*_ is the fractional tissue volume of CSF as determined by the *Seg* module.
- Finally, Osprey derives fully tissue-and-relaxation-corrected molal concentration estimates according to the *Gasparovic* method (Gasparovic et al., 2006), according to

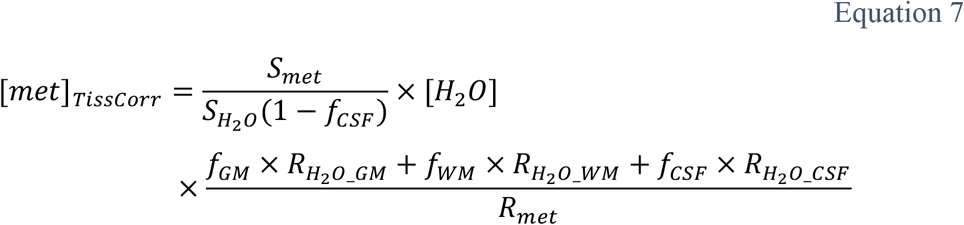

Here, *f*_*GM*_, *f*_*WM*_ and *f*_*CSF*_ are the molal water fractions for GM, WM and CSF, which are derived from the volume fractions according to

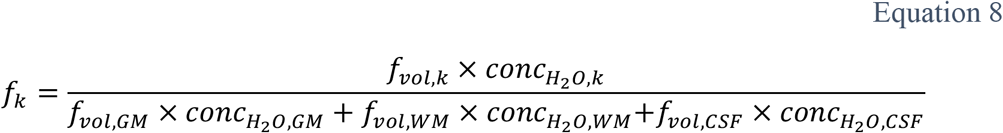

with the relative water densities 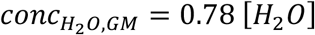, 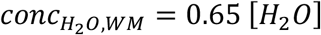, and 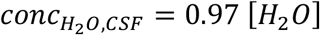. Tissue-specific relaxation corrections are calculated according to 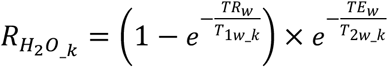 and 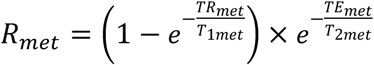. Relaxation times for metabolites and water at 3T field strength were adapted from several widely used references (Edden et al., 2012; Mlynárik et al., 2001; Puts et al., 2013; Wansapura et al., 1999; Wyss et al., 2018). For GABA-edited spectra, Osprey calculates an additional ‘alpha correction’ metric (Harris et al., 2015) that normalizes for the fact that GABA levels are higher in gray matter than in white matter (Jensen et al., 2005).

Osprey saves all available quantitative results for the entire job as comma-separated value (CSV) tables to a subfolder “QuantifyResults” in the output folder. This easily accessible format provides a direct interface to external third-party software for subsequent statistical analysis and visualization (e.g. R, SPSS).

### 2.8 GUI

While all Osprey analysis steps can be carried out with simple MATLAB console commands, the Osprey GUI serves as the central hub for data visualization and quality assessment. Once a job file is loaded, all analysis steps can be triggered with dedicated GUI buttons, and the results from each step can be viewed on separate tabs, for each sub-spectrum from each dataset in the job. The ‘Load’ tab displays the pre-aligned, pre-averaged data, while the ‘Process’ tab shows the aligned individual transients, the final averaged spectra that are passed on to the *Fit* module, as well as information about the frequency history of the experiment pre- and post-alignment, and the basic quality assessment metrics (linewidth and SNR). The ‘Fit’ tab shows the spectrum, the complete fit, residual, baseline, and individual basis function contributions. The ‘Coreg/Seg’ tabs visualizes the results of voxel coregistration and segmentation, and the ‘Quantify’ tab features tables with all available quantitative outcome measures.

The additional ‘Overview’ tab provides useful summary visualizations of batched jobs with many datasets. It provides visualization of mean spectra with overlaid ribbon plots of the standard deviation; mean fit, residual, and baseline; raincloud plots (Allen et al., 2019; Whitaker et al., 2019) of quantitative results for quick assessment of the population distributions of metabolite estimates; and interactive display of correlation plots between metabolite estimates. If the job file was specified in CSV format and included assignment of each dataset to a group (e.g. patients or control subjects), the data in each raincloud and correlation plot is separated by the group variable.

### 2.9 Demonstration

To demonstrate the versatility of the processing and modelling capabilities of Osprey for various acquisition techniques, several single-voxel MRS datasets from the Big GABA dataset (Mikkelsen et al., 2019, 2017) were loaded, processed, and modelled (see section S5 of the Supplementary Materials for details on how to obtain the data from the Big GABA online repository).

Twelve PRESS datasets acquired on a 3.0T GE scanner (GE Healthcare, Milwaukee, United States) were selected. Parameters included: TR/TE = 2000/35 ms; 32 averages; 30 × 30 × 30 mm^3^ voxel in midline parietal cortex; 5 kHz bandwidth with 4096 data points. For comparison and validation, the PRESS datasets were also modelled with LCModel and Tarquin, using default settings, and the same basis functions as for the Osprey analysis. Estimates of total NAA (tNAA = NAA + NAAG), total choline (tCho = GPC + PCh), myo-inositol (Ins) and Glx (glutamate + glutamine) were calculated with respect to total creatine (tCr = Cr + PCr). Agreement of metabolite estimates across software tools was determined by calculating Pearson correlation coefficients separately for each metabolite and each pair of software tools.

Further, eleven GABA-edited MEGA-PRESS datasets from the Big GABA repository, acquired on a 3.0T Philips scanner (Philips Healthcare, Best, The Netherlands), were loaded, processed, and modeled. Parameters that differed from the PRESS parameters included: TE = 68 ms; 320 averages; 15-ms editing pulses applied at 1.9 ppm (edit-ON) and 7.5 ppm (edit-OFF).

Finally, eight GABA/GSH-edited HERMES datasets, acquired on a 3.0T Philips scanner (Philips Healthcare, Best, The Netherlands), were loaded, processed, and modelled. Parameters that differed from the MEGA-PRESS parameters included: TE = 80 ms; 20-ms editing pulses applied in the GABA/GSH HERMES scheme (Saleh et al., 2016).

## 3. Results

All twelve PRESS datasets were successfully loaded and processed in Osprey, and are plotted in Figure 2A. The mean of these processed spectra (and ± one standard deviation range) is shown in Figure 2B, along with the mean of the model spectra and the mean modelling residual. NAA SNR was 150 ± 24 and NAA linewidth was 7.8 ± 1.2 Hz. The results of quantification of these spectra are summarized in the following. tCr ratios of tNAA, tCho, mI, and Glx were 1.46 ± 0.07, 0.18 ± 0.01, 0.75 ± 0.07, and 1.38 ± 0.07, respectively. Water-scaled estimates of tNAA, tCho, mI and Glx, for example, were 19.40 ± 1.27 i.u., 2.42 ± 0.22 i.u., 8.88 ± 0.96 i.u., and 18.42 ± 1.34 i.u., respectively.

**Figure 2:**
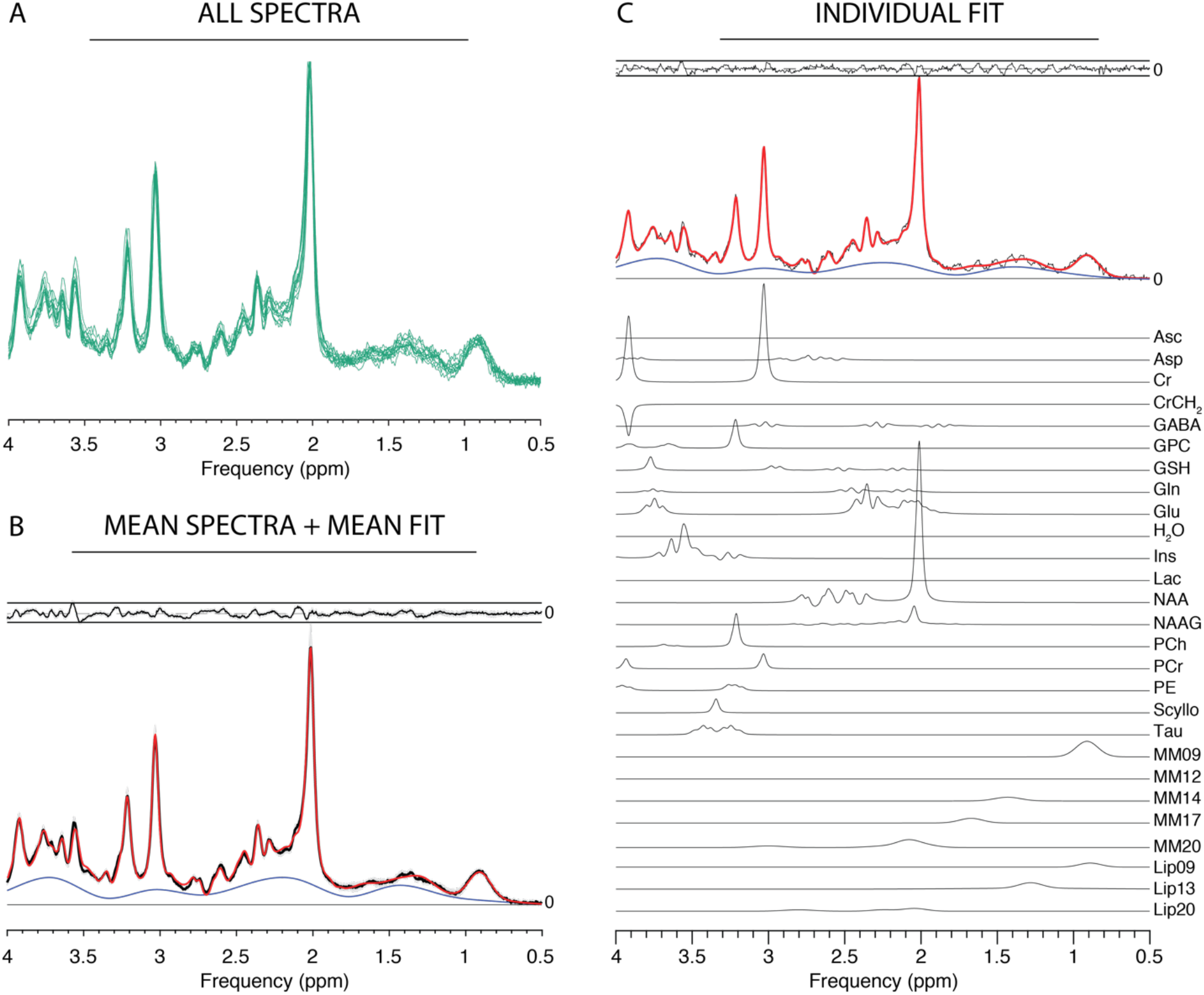
Results from the Osprey processing and modelling of PRESS data. (A) Individual spectra; (B) mean spectra (black) +/− SD (gray ribbons); mean fit (red), mean residual (above). (C) Example fit with contributions from individual metabolites (Asc ascorbate; Asp aspartate; Cr creatine; GABA γ-aminobutyric acid; GPC glycerophosphocholine; GSH glutathione; Gln glutamine; Glu glutamate; Ins myo-inositol; Lac lactate; NAA N-acetylaspartate; NAAG N-acetylaspartylglutamate; PCh phosphocholine; PCr phosphocreatine; PE phosphoethanolamine; Scyllo scyllo-inositol; Tau taurine).

At group level, tNAA/tCr estimates agree well between the three tools, while Tarquin estimates higher tCho/tCr and Glx/tCr, and lower Ins/tCr values than LCModel and Osprey (Figure 3A). Figure 3B shows cross-tool correlation results across the twelve datasets for the four metabolite-to-creatine ratios. Osprey estimates of tNAA/tCr, tCho/tCr and Ins/tCr show significant positive correlations of moderate strength with LCModel estimates. Likewise, tNAA/tCr and Ins/tCr estimates from Osprey and Tarquin correlate positively and moderately strong. In contrast, LCModel and Tarquin correlations do not reach significance.

**Figure 3:**
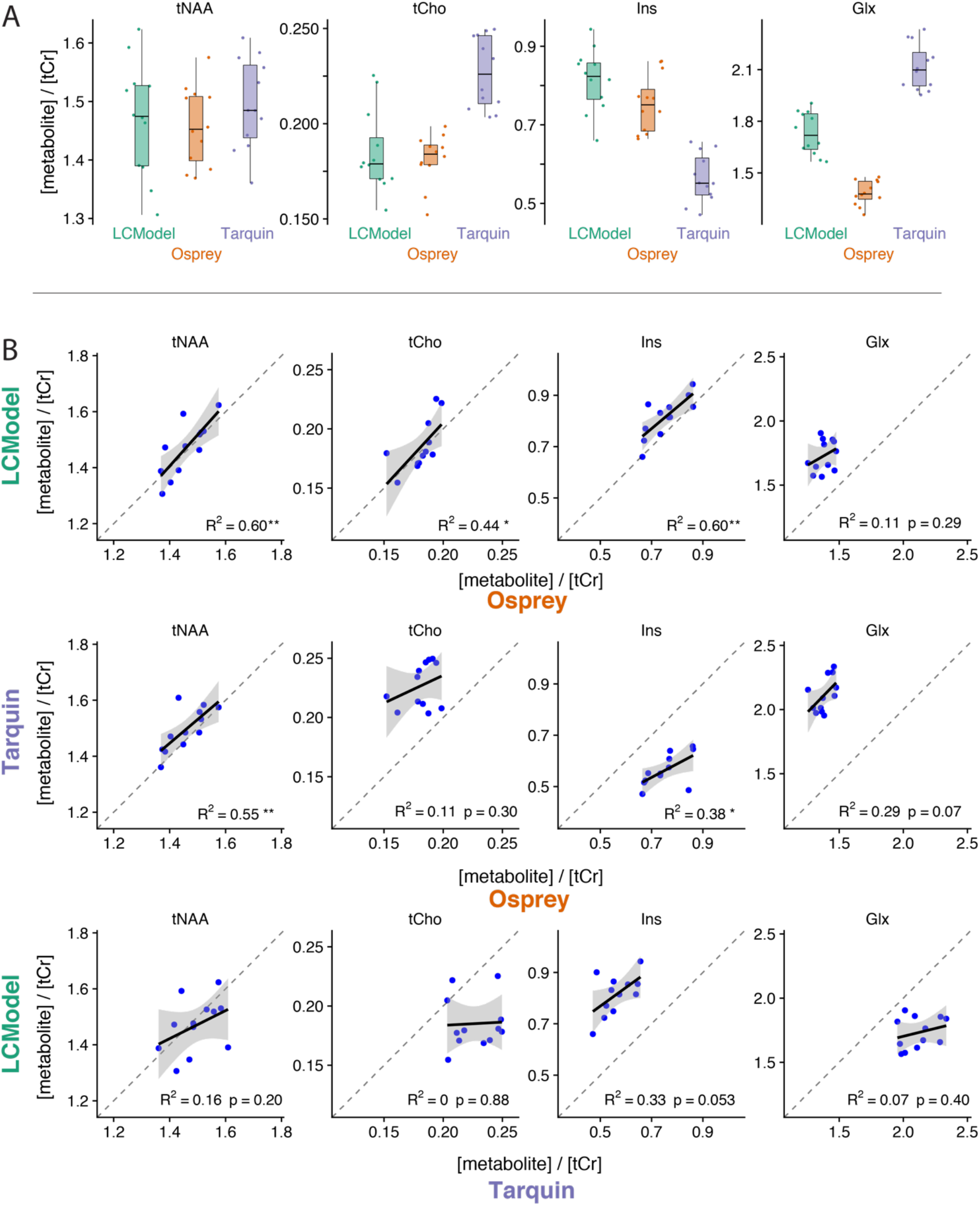
Comparison of metabolite estimates (relative to tCr) for 12 PRESS datasets. (A) Distributions for each metabolite and software; (B) scatterplots for each metabolite and pair of software (upper row: LCModel vs. Osprey, middle row: Tarquin vs. Osprey, bottom row: LCModel vs. Tarquin). One asterisk indicates p < 0.05; two asterisks indicate p < 0.01.

GABA-edited MEGA-PRESS datasets were successfully loaded, processed and modelled in ‘Separate’ mode, as summarized in Figure 4A and B. NAA SNR was 214 ± 32, while the NAA linewidth was 5.1 ± 0.6 Hz. GABA levels were quantified as 0.27 ± 0.07 (tCr ratio) and 1.67 ± 0.31 i.u. (water-scaled), respectively.

**Figure 4:**
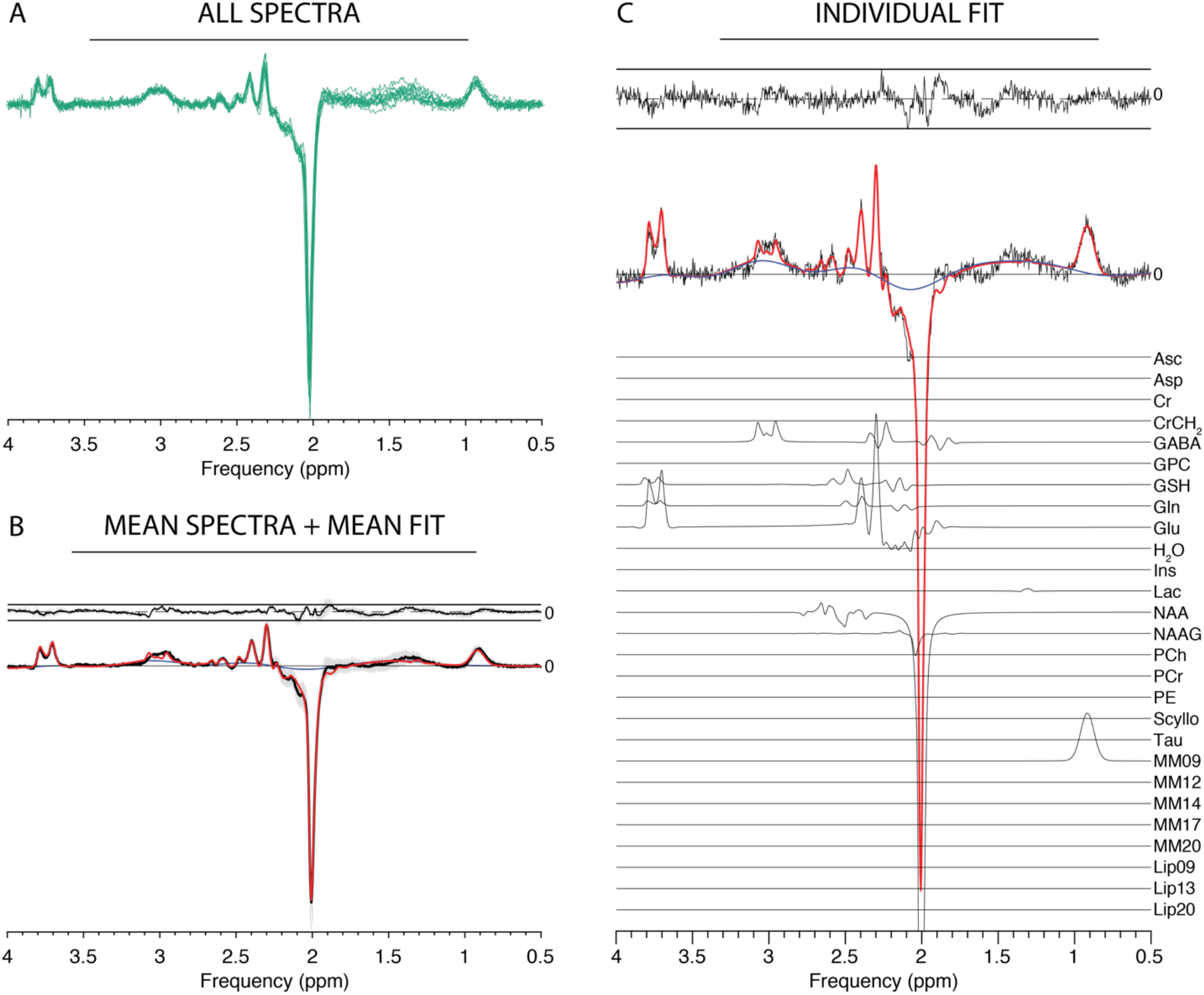
Results from the Osprey processing and modelling of GABA-edited MEGA-PRESS data. (A) Individual spectra; (B) mean spectra (black) +/− SD (gray ribbons); mean fit (red), mean residual (above). (C) Example fit with contributions from individual metabolites.

Similarly, the HERMES data are summarized in Figure 5. NAA SNR was 215 ± 24 with a NAA linewidth of 5.3 ± 0.5 Hz. GABA and GSH levels were estimated as 0.19 ± 0.04 and 0.19 ± 0.03 (tCr ratios), and 2.03 ± 0.28 i.u. and 1.94 ± 0.35 i.u. (water-scaled), respectively.

**Figure 5:**
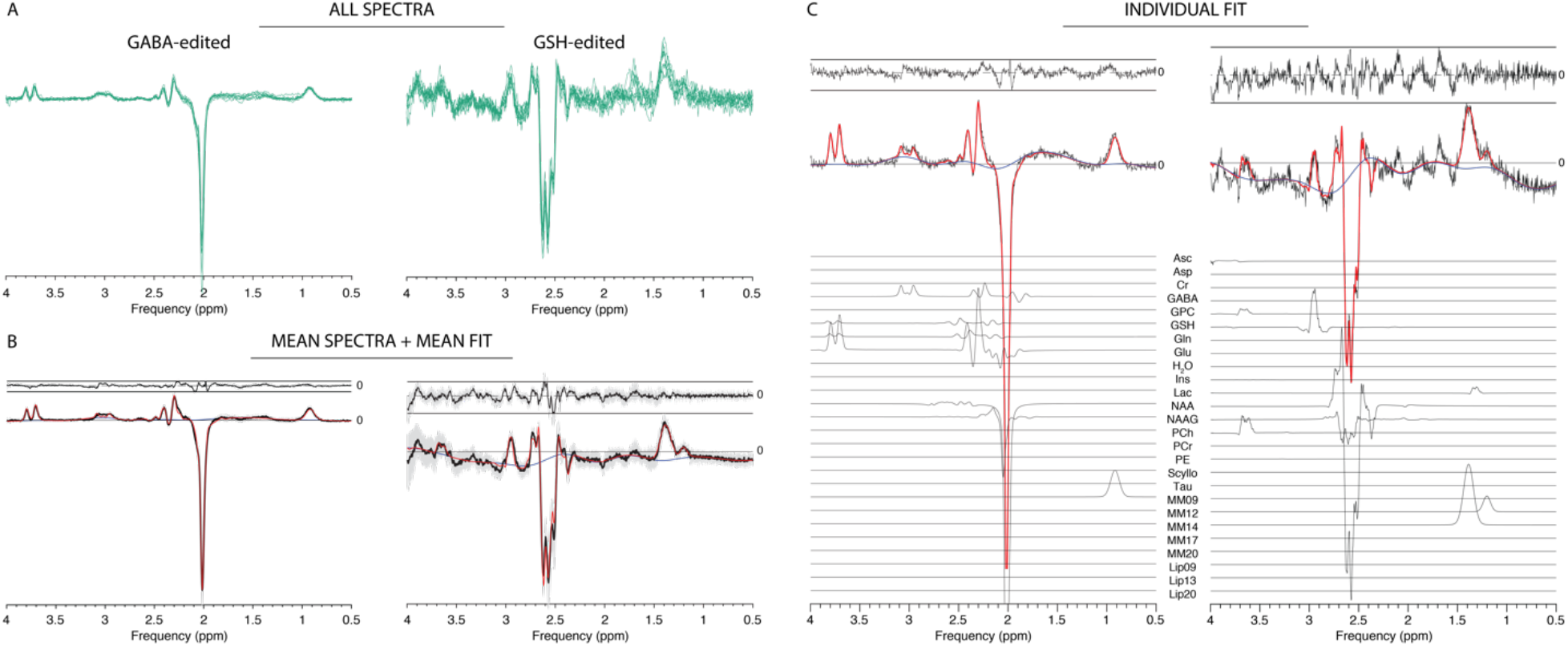
Results from the Osprey processing and separate-mode modelling of GABA/GSH-edited HERMES data. (A) Individual spectra; (B) mean spectra (black) +/− SD (gray ribbons); mean fit (red), mean residual (above). (C) Example fit with contributions from individual metabolites.

The figures show individual spectra overlaid (green lines) in panel A, demonstrating consistent high-quality data resulting from the Osprey processing pipeline. Panels B display mean spectra (black solid lines), the standard deviation of the spectra (gray ribbon plots), mean model fits (red) and mean residuals (above the spectra) across all datasets. Panels C show representative linear-combination modelling. The fits approximate the data well with a relatively smooth baseline.

The structure of the GUI is shown in Figure 6. The workflow buttons corresponding to the different Osprey modules are in the left column, as well as a list of loaded datasets from which the user selects the dataset to be displayed. The tabs above the data display panels are used to switch between the analysis stages, and the tabs below correspond to different sub-spectra (here A, B, C, D, sum and difference spectra for HERMES data, along with water reference data).

**Figure 6:**
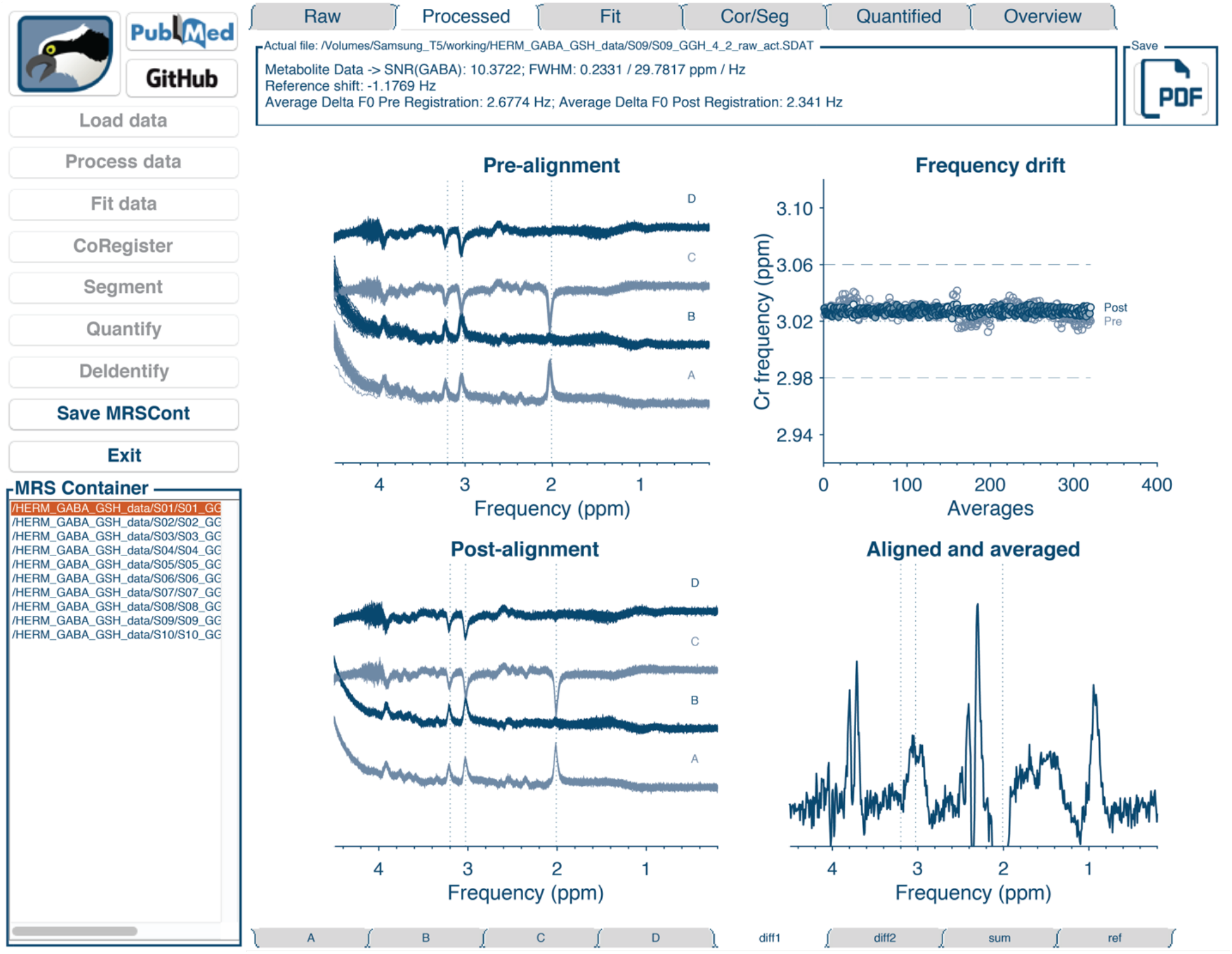
The Osprey GUI with workflow buttons (left), list of datasets (bottom left), analysis stage selection tabs (top row), sub-spectrum selection tabs (bottom row), and data display panel. The figure shows a GABA-GSH-edited HERMES dataset, with sub-experiments A, B, C, and D.

Figure 7 shows the GUI data display panels corresponding to the various stages of analysis of a single HERMES dataset, exemplifying the visualization of spectral-editing sub-experiments.

**Figure 7:**
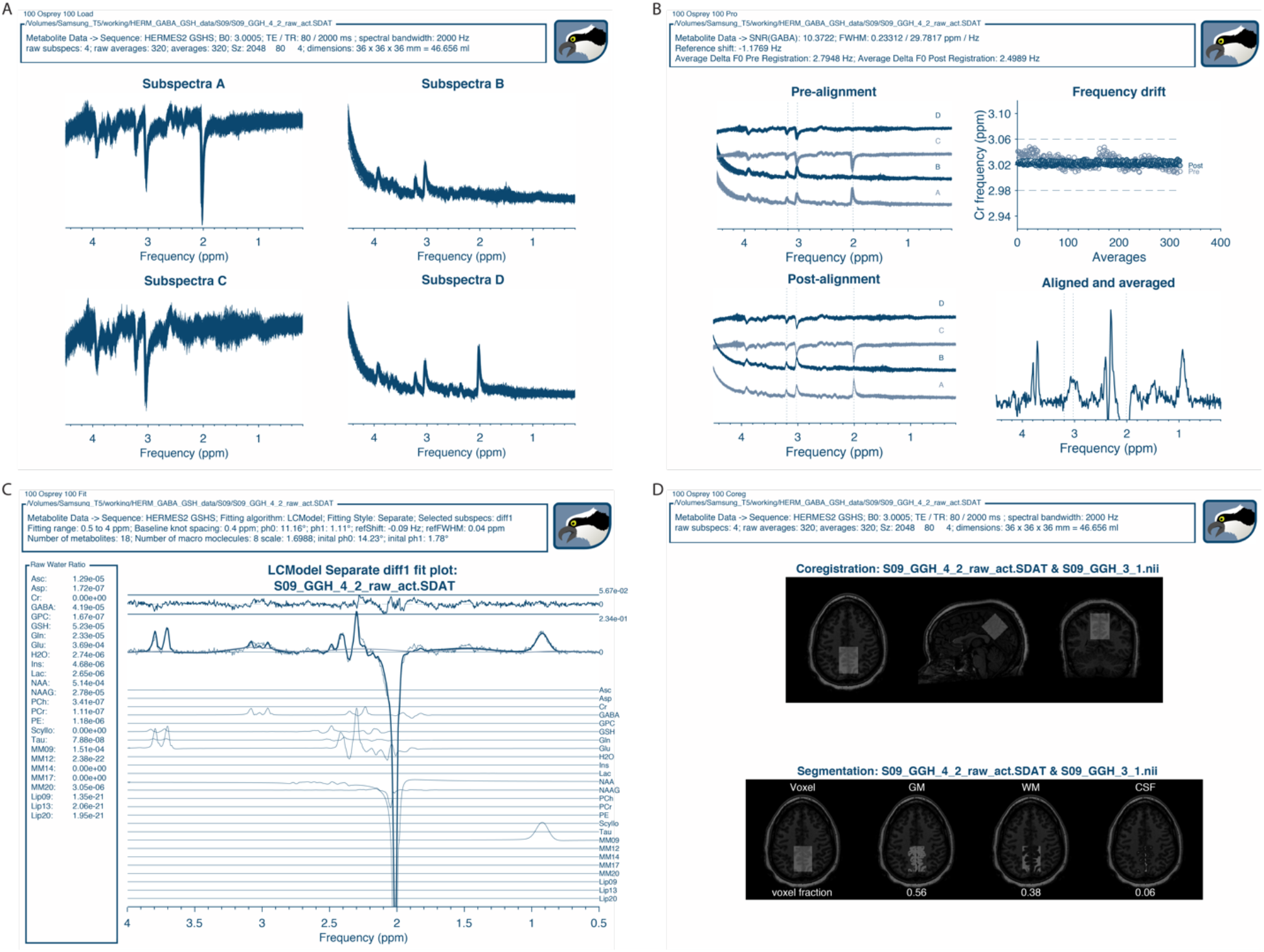
Osprey visualization of the analysis of a HERMES dataset. (A) ‘Load’ visualization; (B) ‘Proc’ visualization with individual transients and frequency drift plot before and after spectral alignment; (C) ‘Fit’ visualization of the GABA-edited HERMES difference spectrum; (D) ‘Coreg’ and ‘Seg’ visualization of voxel coregistration and segmentation, including display of tissue fractions.

## 4. Discussion

The magnetic resonance spectrum of the human brain is rich with biochemical information, but extracting that information is a challenging task due to the overlapped nature of the metabolite spectra and the broad in-vivo linewidth. Quantitative MRS measurement outcomes are known to vary considerably depending on field strength, scanner vendor, localization technique, acquisition parameters, and choice of data processing and quantification practices. Recently, the MRS community has led efforts to converge towards standardized data acquisition (Deelchand et al., 2019a; Öz et al., 2020; Saleh et al., 2019; Wilson et al., 2019). In contrast, consensus on the processing and analysis of data has been slower to emerge.

The most commonly used strategy for quantitative analysis of MR spectra is linear-combination modelling. As a result of the methodological diversity and lack of standardized processing pipelines, most researchers have developed their own code to prepare their data for third-party quantification software. This practice is problematic for a number of reasons: a) methodological heterogeneity and opacity diminish comparability and reproducibility of quantitative MRS studies; b) benchmarking and subsequent adaptation of methodological progress is considerably slowed down; c) researchers new to the field experience a high-level entry threshold; d) strong dependency on engagement, support, and funding situation of third-party software developers leaves the community vulnerable.

Our new toolkit ‘Osprey’ seeks to address these issues by providing the community with a freely available open-source environment that unifies all steps of modern MRS data analysis – processing, modelling, quantification – into a common framework. Osprey is designed to: a) reduce methodological heterogeneity with its built-in standardized processing pipeline, modelling and quantification routines. These can serve as a substrate and starting point for the development of advanced data analysis methods, such as coil-combination or spectral alignment routines; b) accelerate the benchmarking, critical evaluation and finally integration of improved methods into the modular workflow; c) be immediately deployed by novice MRS users who seek to obtain quantitative results from MRS data using a single software solution, and who lack the resources to develop analysis code of their own; d) reduce the dependency of the MRS community on continued development of third-party linear-combination modelling software. Instead, the model code is directly accessible, modifiable, and exchangeable, allowing the research community to study the factors influencing results of linear-combination modelling, and compare or improve modelling algorithms.

Across the demonstration sample of PRESS datasets, Osprey estimates of the major metabolites are comparable to estimates from LCModel and Tarquin, while the agreement between LCModel and Tarquin was weaker than expected. Interestingly, performance and agreement of different linear-combination algorithm have not been systematically assessed on a larger scale. The influence of critical experimental parameters and analysis settings that critically affect performance and results of different modeling algorithms also remain under-studied, for example the influential ‘DNKTMN’ parameter that determines the degree of spline baseline flexibility in LCModel (Bhogal et al., 2017; Marjanska and Terpstra, 2019), or interactions of metabolite estimates with linewidth or SNR (Near, 2014). For difference-editing experiments, there is currently no consensus on optimal linear-combination strategies. In particular, co-edited macromolecular signals are difficult to simulate appropriately, since the spin systems of the contributing molecules are poorly characterized. Currently, co-edited signals representing the MM09, MM12 and MM14 resonances are therefore empirically parametrized in Osprey as Gaussian-shaped. Using Osprey, advanced modeling strategies for edited experiments can be explored, most notably for GABA editing experiments with unresolved macromolecular contamination issues, e.g. whether to include macromolecular and homocarnosine basis functions (Deelchand et al., 2019b), whether to impose soft constraints on co-edited macromolecules (Murdoch and Dydak, 2011), whether to increase baseline stiffness, whether to constrain the model by incorporating fit information from the sum spectrum (Oeltzschner et al., 2019a, 2019b), how to perform appropriate GABA-specific tissue correction strategies (Harris et al., 2015) etc. Osprey facilitates methodological investigations like these through its job system that allows many datasets to be batch-processed by modifying a single text file. While large-scale repositories of MRS data are still rare, projects like Big GABA (https://www.nitrc.org/projects/biggaba/) already provide publicly available datasets that can be easily deployed to benchmark the performance of analysis methods with high statistical power.

Many other MRI modalities suffer from susceptibility to ‘processing bias’, and their communities have developed and adopted de-facto-standardized data processing and analysis toolboxes. Notable examples are, among others, SPM (Friston, 2007) and FSL (Jenkinson et al., 2012) for fMRI analysis, BART (Uecker et al., 2014) for parallel imaging reconstruction, and the Spinal Cord Toolbox (De Leener et al., 2017). In contrast, no such common framework currently exists for MRS, but rather a diverse field of tools mostly dedicated to modelling or visualization. LCModel continues to be the most widely used spectral analysis tool, despite its current cost of 13,300 USD, restrictive licensing, limited ongoing development by a single software engineer, and lack of built-in pre-processing functions. jMRUI offers basic functions to manipulate spectra and a variety of time-domain modelling algorithms (AMARES, QUEST, AQSES), but requires a high degree of user interaction and expertise, making it less suitable for novice researchers and reproducible processing of large datasets. Tarquin supports automated processing, but, like jMRUI, requires pre-processed spectra to model, and leaves a lot of freedom in choosing modelling options to the user. Gannet is an open-source toolbox with a similar all-inclusive workflow as Osprey, but is limited to simple peak integration of spectral-edited data. The MATLAB toolbox OXSA (Purvis et al., 2017) is conceptually similar to Osprey, but currently limited to processing DICOM data, and implements the AMARES algorithm (Vanhamme et al., 1997), which fits spectra in the time-domain using series of singlets with user-imposed prior knowledge, an approach notably different from linear-combination modelling of simulated metabolite basis functions. SIVIC (Crane et al., 2013) does not offer data pre-processing or linear-combination modelling at all, and is primarily dedicated to visualizing MRSI data and interfacing with radiological PAC systems. INSPECTOR (Juchem, 2018) offers automated analysis, but has not been peer-reviewed and is only distributed as closed-source obfuscated MATLAB executable code, as is the more interactive MRspa (Center for Magnetic Resonance Research, University of Minnesota, 2018).

All these existing software solutions either run as compiled executables or closed-source MATLAB applications, and while the source code is publicly available for some (Tarquin, SIVIC, Vespa), community-sourced modifications are either impossible, or may require substantial modifications downstream and local recompiling. In contrast, the entire Osprey source code (written in MATLAB) is publicly available in a single repository at https://github.com/schorschinho/osprey. At the time of writing, Osprey functions are primarily tailored to processing, modelling, and quantifying data from widely used single-voxel ^1^H MRS sequences designed to detect common metabolites in the human brain *in-vivo*. The modularity of the pipeline allows developers and users to implement analogous workflows for data acquired with different localization techniques, at different field strength, from other nuclei and body parts, animals, or phantoms, simply by branching out the workflow using ‘if’ statements and flags. Our group has previously disseminated the open-source processing toolkit Gannet (Edden et al., 2014), which has since been continuously developed in close interaction with its user base. We are confident that this pool of collaborators (and potential code contributors) will provide a solid foundation for continued development support for Osprey in the future.

## 5. Conclusions

Osprey is a new, open-source software environment for the pre-processing, linear-combination modelling, quantification and visualization of magnetic resonance spectroscopy data. It is hoped that the availability of such a tool will improve the standardization and accessibility of MRS data processing, while enabling further investigation and rapid adoption of new methodology.

## Acknowledgements

This work was supported by NIH grants K99AG062230, R21AG060245, R01EB016089, R01EB023963, and P41EB015909. None of the authors have a competing interest to declare.

