## Supplementary Material for "Osprey: Open-Source Processing, Reconstruction & Estimation of Magnetic Resonance Spectroscopy Data"

**S1. Details and commands of the Osprey analysis workflow**

1. At the beginning of every Osprey analysis, a fresh MRSCont data container with a job (defined in the file jobFile, see “Detailed structure of an Osprey job file” below) is initialized using the following command:

[MRSCont] = OspreyJob(jobFile);

Upon execution, the *OspreyJob* command initializes a MATLAB structure array that serves as the superstructure for all settings and data that have been previously specified in the job file. All subsequent modules of the Osprey pipeline act solely on the superstructure created by the *OspreyJob* module. The name of the superstructure variable (MRSCont in this example) is the only argument to be passed on when calling other functions along the Osprey workflow.

The job file can specify an additional file (in CSV format), which allows the specification of external statistical variables for each dataset, such as subject age, diagnostic classifiers, or behavioral measures.

2. After loading a job file, the data loading process can be initiated by executing the command:

[MRSCont] = OspreyLoad(MRSCont);

Upon calling the *OspreyLoad* command, Osprey parses the filename endings of the MRS data files to determine the correct file format. The appropriate loading functions then extract all relevant information from the headers to load the raw spectroscopic data into the Osprey superstructure. Aside from the FIDs, the *Load* module saves the receiver bandwidth, repetition and echo times, number of averages, number of data points, transmitter frequency (or magnetic field strength), as well as information about voxel dimensions and positions.

The *Load* module also combines the signals from multi-channel receiver coils by determining the channel-specific phase and weighting each channel with the ratio of the signal to the square of the noise. Assuming uncorrelated noise, this procedure has been shown to yield the optimal signal-to-noise ratio (Hall et al., 2014). If lineshape reference or short-TE reference data are provided, they are used instead of the metabolite data to determine the phasing and weighting parameters.

The coil-combined (but un-aligned and un-averaged) data is stored in FID-A data structure arrays within the Osprey superstructure for further processing.

3. Once the raw data is loaded, the next function *OspreyProcess* initiates a series of automatic processing steps, using the command:

[MRSCont] = OspreyProcess(MRSCont);

This step performs all state-of-the-art MRS post-processing techniques to ensure optimal spectral quality, including eddy-current correction, frequency-and-phase correction, removal of residual water signal, baseline correction, and automated phasing and frequency referencing. In addition, several basic quality metrics (linewidth, SNR) are determined.

The *OspreyProcess* module is also responsible for handling all aspects of data processing that are specific to J-difference-edited ("spectral editing") experiments, such as MEGA, HERMES, and HERCULES. These additional steps include alignment of sub-spectra to reduce subtraction artefacts, as well as the calculation of (Hadamard-encoded) difference spectra.

If appropriate flags are being set in the job file (see “Detailed structure of an Osprey job file”), Osprey saves the fully processed spectra in formats readable to LCModel, jMRUI, Tarquin, etc. For spectral editing data, separate files are created for the difference and sum spectra, as well as for each sub-experiment (edit-ON/OFF for MEGA, A/B/C/D for HERMES/HERCULES). Additional files are created for the lineshape and short-TE water data, if available. If water reference data are acquired with spectral editing sequences, the subspectra are added and saved as a single file.

4. The linear-combination modelling process is started using the command

[MRSCont] = OspreyFit(MRSCont);

*OspreyFit* performs linear-combination modeling of the processed spectra. The Osprey fitting algorithm uses metabolite basis functions for the specific MRS sequence, which are calculated from 2-D spatially resolved full density-matrix simulations, using the real sequence timings and pulse waveforms wherever possible. Osprey basis sets can be generated by users or upon request by the authors, and added to the repository. Basis sets can be created with a single command-line function (fit_makeBasis.m) from a set of spectra that have been simulated with FID-A. Alternatively, existing LCModel basis sets (.BASIS) can be imported into Osprey format using another single command-line function (io_LCMBasis.m).

For J-difference-edited experiments, the user can choose between two fitting styles in the job file. When the ‘Concatenated’ fitting style is selected, all difference spectra and the sum spectrum are modeled simultaneously. In this case, the amplitude, lineshape, phase and linebroadening parameters are shared between the models for each sub-spectrum, while each sub-spectrum maintains its own baseline parameters and is allowed an additional small frequency shift to account for small inconsistencies between sub-spectra. The simultaneous modelling approach incorporates all available spectral information to constrain the model in a way that is most consistent with the data, thereby reducing the variability of the quantification results compared to unconstrained separate modelling (Oeltzschner et al., 2019a). When the ‘Separate’ fitting style is selected, each difference spectrum is modeled separately – one for MEGA-edited data, two (or more) for Hadamard-edited data like HERMES and HERCULES. In this case, the editing-off spectrum (for MEGA) or the sum spectrum (for HERMES/HERCULES) is also separately modeled.

5. If T1-weighted anatomical images in NIfTI format have been provided in the job file, co-registration can be initiated with the command:

[MRSCont] = OspreyCoreg(MRSCont);

This function parses voxel geometry information that was extracted from the MRS data headers by OspreyLoad. While the definitions of the voxel orientation parameters differ between vendors, in each case, they uniquely define the dimensions and positioning of the voxel in scanner space. SPM12 (<https://www.fil.ion.ucl.ac.uk/spm/software/spm12/>) NIfTI handling functions are then called to create an SPM image volume containing a binary voxel mask. The voxel masks are saved in NIfTI format using SPM volume processing tools in a sub-directory (/VoxelMasks/) of the output folder specified in the job file.

6. Once the coregistration module has been completed, tissue segmentation can be called using:

[MRSCont] = OspreySeg(MRSCont);

This function uses the SPM12 segmentation function to segment the structural images into gray matter (GM), white matter (WM), and cerebrospinal fluid (CSF). The fractional tissue volumes are subsequently used for water-scaled metabolite quantification.

7. Finally, the quantification module can be invoked with the command:

[MRSCont] = OspreyQuantify(MRSCont);

This module calculates various quantitative outputs, depending on the available modelling parameters that have been determined during the *Fit* process, including ratios to total creatine, raw water-scaled estimates, CSF-corrected estimates, and estimates fully corrected for tissue water content and relaxation effects.

**S2. Detailed structure of an Osprey job file**

The Osprey job file is the only point of direct contact between the user and the analysis, ensuring that all processing, modeling, and quantification steps are performed in an operator-independent, reproducible way.

The user specifies all necessary information through a single text file, or alternatively in the form of a CSV-formatted table. The following pieces of information are required for a valid job file:

1. Specify full paths to the raw data
   a. Metabolite (water-suppressed) data
   b. OPTIONAL: Lineshape reference (water-unsuppressed) data – has to be acquired with the same sequence parameters as the metabolite data
   c. OPTIONAL: Short-TE water reference (water-unsuppressed) data – for quantification purposes with reduced T2 weighting only
   d. OPTIONAL: Structural images (*.nii)
2. Sequence information
   a. Type (conventional/unedited, MEGA, HERMES, HERCULES)
   b. OPTIONAL (when edited data is provided): Spectral editing target metabolite(s)
3. Data export
   a. OPTIONAL: Save processed spectra in LCModel format?
   b. OPTIONAL: Save processed spectra in jMRUI format?
   c. OPTIONAL: Save processed spectra in vendor-specific native format?
4. Modeling options
   a. Model the spectra in multiplexed datasets (e.g. edited data) separately or simultaneously?
   b. Fit range in the frequency domain
   c. Baseline knot spacing
   d. Add macromolecule and lipid basis functions?
5. Specify full path to an output folder
   a. Specify folder where the Osprey data container, exported third-party format files, and result tables are saved
6. Additional dataset-specific information
   a. OPTIONAL: Specify full path to a CSV file containing subject-specific information such as age, diagnostic group, behavioral measures, etc.

**S3. Definitions of macromolecule and lipid basis functions**

For unedited data, macromolecule and lipid basis functions are created in analogy to the definitions used in LCModel and Tarquin (Table S1):

| *Name* | *Frequencies [ppm]* | *FWHM [ppm]* | *Amplitude* |
| --- | --- | --- | --- |
| MM09 | 0.91 | 0.14 | 3.00 |
| MM12 | 1.21 | 0.15 | 2.00 |
| MM14 | 1.43 | 0.17 | 2.00 |
| MM17 | 1.67 | 0.15 | 0.20 |
| MM20 | 2.08 | 0.15 | 1.33 |
|  | 2.25 | 0.20 | 0.33 |
|  | 1.95 | 0.15 | 0.33 |
|  | 3.00 | 0.20 | 0.40 |
| Lip09 | 0.89 | 0.14 | 3.00 |
| Lip13a | 1.28 | 0.15 | 2.00 |
| Lip13b | 1.28 | 0.089 | 2.00 |
| Lip20 | 2.04 | 0.15 | 1.33 |
|  | 2.25 | 0.15 | 0.67 |
|  | 2.80 | 0.20 | 0.87 |

Table S1: Default properties of the macromolecule and lipid basis functions included in Osprey basis sets, mimicking the approach of LCModel (section 11.7 of the LCModel manual). Amplitude values are scaled relative to the 3.02 ppm Cr CH_3_ singlet in the basis set, which has amplitude 3.

Table S2 summarizes the parametrization of co-edited macromolecule signals in edited difference spectra. These parameters have been derived empirically on the datasets used in this manuscript, and may require further optimization. For GABA-edited data (MEGA-PRESS and HERMES), the basis functions include a resonance for the 0.91 ppm macromolecule (MM09) signal. For GSH-edited HERMES data, resonances for the 1.2 and 1.4 ppm macromolecules (MM12, MM14) are included:

| *Name* | *Frequencies [ppm]* | *FWHM [ppm]* | *Amplitude* |
| --- | --- | --- | --- |
| MM09 | 0.915 | 0.085 | 3.00 |
| MM12 | 1.20 | 0.07 | 2.00 |
| MM14 | 1.385 | 0.095 | 2.00 |

Table S2: Default properties of the macromolecule and lipid basis functions included in Osprey basis sets for GABA- and GSH-edited data. Amplitude values are scaled relative to the 3.02 ppm Cr CH_3_ singlet in the edit-OFF basis function, which has amplitude 3.

**S4. Modelling constraints**

Default hard constraints on non-linear parameters and weak soft constraints on macromolecule and lipid amplitudes are imposed to stabilize the solution, and are defined as they are in LCModel and Tarquin (Table S3). Here, $\varphi_{0}$ represents the global zero-order phase correction; $\varphi_{1}\left( \nu\right)$ the global first-order (linear) phase correction, $\gamma$ the Gaussian linebroadening, $\alpha_{m}$ the individual Lorentzian linebroadenings and $\omega_{m}$ ($m=1,\ldots)$ the individual frequency shifts for all $N_{M}$ basis functions:

| **Hard constraints** | *Parameter* | *Lower bound* | *Upper bound* | **Weak soft constraints** | *Metab 1* | *Metab 2* | *Ratio* |
| --- | --- | --- | --- | --- | --- | --- | --- |
|  | $\varphi_{0}$ | $-359^{\circ}$ | $359^{\circ}$ |  | NAAG | NAA | 0.15 |
|  | $\varphi_{1}$ | $-10^{\circ}/ppm$ | $10^{\circ}/ppm$ |  | Lip09 | Lip13 | 0.267 |
|  | $\gamma$ | $0 Hz$ | $\sqrt{5000} Hz$ |  | Lip20 | Lip13 | 0.15 |
|  | $\alpha_{m}$ (metabolites) | $0 Hz$ | $10 Hz$ |  | MM20 | MM09 | 1.5 |
|  | $\alpha_{m}$ (MM/lipids) | $0 Hz$ | $100 Hz$ |  | MM12 | MM09 | 0.3 |
|  | $\omega_{m}$ (metabolites) | $-0.03 ppm$ | $0.03 ppm$ |  | MM14 | MM09 | 0.75 |
|  | $\omega_{m}$ (MM/lipids) | $-0.05 ppm$ | $0.05 ppm$ |  | MM17 | MM09 | 0.375 |

*Table S3: Hard constraints on fitting parameters, and weak soft constraints on amplitude ratios.*

**S5. Datasets included in the analysis**

The PRESS data from site G4 analyzed in the manuscript can be downloaded freely from the Big GABA repository at <https://www.nitrc.org/frs/download.php/11684/G4_P.zip>.

The MEGA-PRESS data from site P3 analyzed in the manuscript can be downloaded freely from the Big GABA repository at <https://www.nitrc.org/frs/download.php/10975/P3_MP.zip>.

The HERMES data analyzed in the manuscript includes the original data from MG Saleh et al, Simultaneous edited MRS of GABA and glutathione (NeuroImage 142:576-582 (2016), <https://doi.org/10.1016/j.neuroimage.2016.07.056>), and can be made available upon request.
